# sc2MeNetDrug: A computational tool to uncover inter-cell signaling targets and identify relevant drugs based on single cell RNA-seq data

**DOI:** 10.1101/2021.11.15.468755

**Authors:** Jiarui Feng, S. Peter Goedegebuure, Amanda Zeng, Ye Bi, Ting Wang, Philip Payne, Li Ding, David DeNardo, William Hawkins, Ryan C. Fields, Fuhai Li

**Affiliations:** Institute for Informatics (I2), Washington University School of Medicine; Department of Computer Science and Engineering; Department of Neurology; Department of Medicine; Department of Surgery; Siteman Cancer Center; Department of Pediatrics, Washington University School of Medicine, Washington University in St. Louis, St. Louis, MO, USA

## Abstract

Single-cell RNA sequencing (scRNA-seq) is a powerful technology to investigate the transcriptional programs in stromal, immune, and disease cells, like tumor cells or neurons within the Alzheimer’s Disease (AD) brain or tumor microenvironment (ME) or niche. Cell-cell communications within ME play important roles in disease progression and immunotherapy response and are novel and critical therapeutic targets. Though many tools of scRNA-seq analysis have been developed to investigate the heterogeneity and sub-populations of cells, few were designed for uncovering cell-cell communications of ME and predicting the potentially effective drugs to inhibit the communications. Moreover, the data analysis processes of discovering signaling communication networks and effective drugs using scRNA-seq data are complex and involve a set of critical analysis processes and external supportive data resources, which are difficult for researchers who have no strong computational background and training in scRNA-seq data analysis. To address these challenges, in this study, we developed a novel open-source computational tool, sc2MeNetDrug (https://fuhaililab.github.io/sc2MeNetDrug/). It was specifically designed using scRNA-seq data to identify cell types within disease MEs, uncover the dysfunctional signaling pathways within individual cell types and interactions among different cell types, and predict effective drugs that can potentially disrupt cell-cell signaling communications. sc2MeNetDrug provided a user-friendly graphical user interface to encapsulate the data analysis modules, which can facilitate the scRNA-seq data-based discovery of novel inter-cell signaling communications and novel therapeutic regimens.

## Introduction

Tumor-stroma communication within the tumor microenvironment (TME) plays an important role in tumor development and responses to both conventional– and immune-based therapies. For example, immunotherapy in pancreatic cancer treatment has not been successful^1^. One possible cause of immunotherapy resistance is the abundance of stromal cells and tumor signaling communications in Pancreatic ductal adenocarcinoma (PDAC) tumor microenvironments^2^. Such immunosuppressive cells include tumor-associated macrophages (TAMs), myeloid-derived suppressor cells (MDSCs), regulatory T cells (Tregs), as well as cancer-associated fibroblasts (CAFs)^2,3,4,5,6,7^. Moreover, CAFs were recently reported to be able to regulate the invasive epithelial-to-mesenchymal transition (EMT) and proliferative (PRO) phenotypes of PDAC^8^. This indicates that stroma-tumor communication in PDAC tumor microenvironments plays a critical role in immunotherapy resistance. Thus, stroma-tumor signaling communications are potential targets to improve drug or immunotherapy response in cancer treatment. The inhibition of signaling communication between TAMs and PDAC cells via the Colony Stimulating Factor 1 (CSF1) (ligand secreted by PDAC) and CSF1R (receptor on TAM) can reprogram TAMs, and the synergistic combination of TAM-tumor signaling inhibition with the immune checkpoint blockade^9^ can improve the immunotherapy response. In another study, the inhibition of signaling communication between CAF and PDAC via CXCL12 (ligand secreted by CAF) and CXCR4 (receptor on PDAC) was shown to improve immunotherapy response^10^. Another example is AD, which is a complex disease with altered inflammation and immune functions in AD brain ME^11–14^. However, the detailed mechanism of how stroma and immune cells like astrocytes and microglia influence the activity of each other and neurons remain unclear. Especially, which signaling pathways and genes are dysfunctional or expressed abnormally. These impede the development of novel drugs and drug combinations for the control and treatment of AD.

Recent advances in single-cell RNA sequencing (scRNA-seq) create a powerful technology to analyze the genetic and functional heterogeneity of stromal and tumor cells (e.g., TAM, CAF and T cells) within tumor microenvironments^15,16,17^. Similarly, studies have generated scRNA-seq data of AD brain samples to investigate the dysfunctions of neurons, astrocytes, microglia cells and other cells in AD brain microenvironments^12–14,18^. Though many tools and studies reported to have discovered the heterogeneity and sub-populations of cells, few studies^19^ have been designed to investigate cell-cell communication using sc-RNAseq data. For example, the CCCExplorer^20,21^ was first developed to uncover the potential tumor and stroma cell communication using microarray and bulk RNA data on a small set of curated ligand-receptor interactions. CellPhoneDB^22^ provided a repository of ligands, receptors, and their interactions using the novel computational ligand-receptor interaction prediction approaches. NicheNet^23^ was the latest software tool that integrates the large set of ligand-receptor interactions from CellPhoneDB, and it supports the pre-analyzed scRNA-seq data. However, the computational modules of inferring the dysfunctional signaling networks, and predicting potentially effective drugs inhibiting the dysfunctional signaling networks and cell-cell communications are not available in these tools.

Specifically, compared with the existing tools, novel computational models and tools that solve the following challenges are in high demand to 1) provide an end-to-end model that can take the raw scRNA-seq data as input, analyze, annotate and display the scRNA-seq data, 2) uncover dysfunctional signaling network within individual cells, and uncover complex signaling communications among multiple stromal and tumor cells; 3) identify effective drugs and drug combinations that disrupt the cell-cell communications, like stroma-tumor, to improve the targeted and immunotherapy response. Moreover, 4) a user-friendly interactive graphical user interface (GUI) is helpful and critical for biomedical researchers because these analyses are highly composite complex and involve a set of computational analysis processes and integration of external supportive data resources that require visualization by non-bioinformatics experts to functionalize the complex data. To resolve the aforementioned challenges, in this study, we developed a novel computational tool: **sc2MeNetDrug** (<u>sc</u>RNA-seq based modeling <u>to</u> discover disease <u>m</u>icro<u>e</u>nvironment signaling communication <u>net</u>works and <u>drug</u>s targeting the cell-cell signaling communications). sc2MeNetDrug provided a user-friendly graphical user interface to encapsulate the data analysis modules, which can facilitate the scRNA-seq data-based discovery of novel inter-cell signaling communications and novel therapeutic regimens. The sc2MeNetDrug, source code, and detailed documentations are publicly available at: https://fuhaililab.github.io/sc2MeNetDrug/.

## Results

### Overview of sc2MeNetDrug

**Fig 1** summarizes the overall pipeline of the **sc2MeNetDrug**. The input of sc2MeNetDrug is the raw counts of genes from single cell RNA-seq (scRNA-seq) data of different experimental conditions or samples, e.g., normal tissues vs disease tissues. The output of the tool includes the annotation of cell types, dysfunctional signaling networks within individual cells, intercellular signaling communications, and drugs that can potentially inhibit dysfunctional signaling pathways and intercellular signaling communications. Specifically, the pipeline can be divided into several parts: First, users need to upload the raw data along with an optional design/group file. Then, the raw data go through preprocessing, dimension reduction, clustering, and cell annotation sequentially to obtain the cell annotation result for each cell (cell annotation results can also be uploaded directly to the application to conduct the rest of analyses). Next, various analysis can be performed based on the interest and requirement, including ***iCSC*** (**i**nter-**<u>c</u>**ell **<u>s</u>**ignaling **<u>c</u>**ommunication discovery) module that uncovers the activated signaling pathways and gene ontology (GO) terms within individual cell types, and uncovers the cell-cell signaling communications within the disease ME and ***dCSC*** (**<u>d</u>**rug prediction for disrupting **<u>c</u>**ell **<u>s</u>**ignaling **<u>c</u>**ommunication) module that identify and predict the potentially effective drugs, based on drug-target and revere gene signature, to disrupt the cell signaling communications. All the data analyses and modeling were designed in the modular format, which can be upgraded or replaced conveniently to select the best practice models. A detailed introduction to the downloading, installation, analysis modules, and examples, as well as the video tutorials for each analysis module, were provided at: https://fuhaililab.github.io/sc2MeNetDrug/. We applied the SC2MeNetDrug model to both a cohort of pancreatic ductal adenocarcinoma (PDAC) and two cohorts of Alzheimer’s disease scRNA-seq data demonstrating the functionality and effectiveness of the tool.

**Figure 1:**
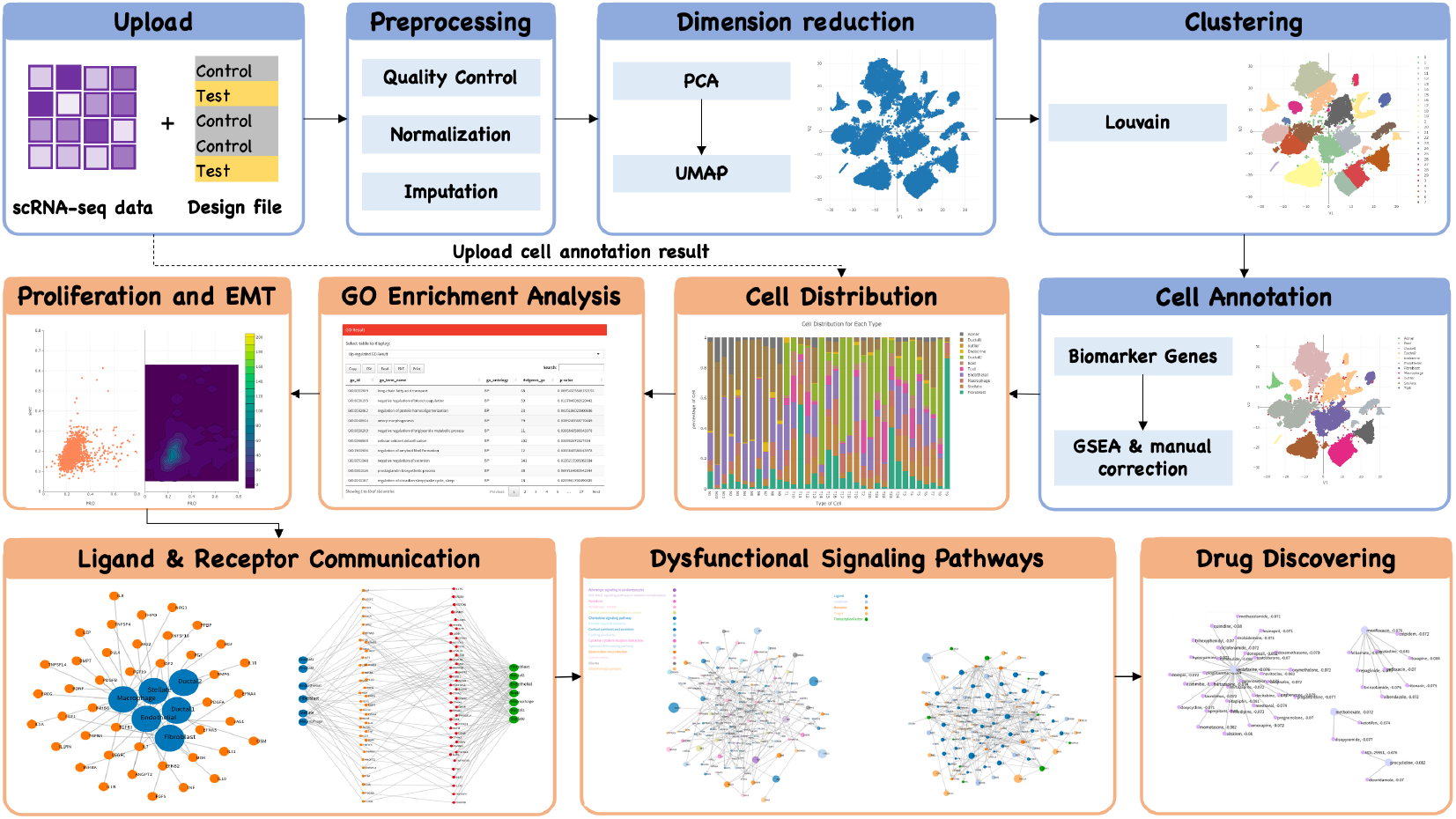
Overview of SC2MeNetDrug. The data analysis pipeline of sc2MeNetDrug can be divided into the following parts: (1) Raw data uploading: Users need to first upload raw-count data along with an optional design file (cell group); (2) Preprocessing: Then, the preprocessing is applied on the raw data to perform the quality control, normalization, and imputation; (3) Dimension reduction: the dimension reduction algorithm applied on the normalized data; (4) Clustering: Cluster cell sample into different group based on dimension-reduced representation for each cell sample; (5) Cell annotation: annotate each cell cluster with the best matching cell type given cell candidate and corresponding marker genes (The cell annotation result can also be uploaded along with raw data to directly perform the following analyses); (6) Cell distribution: Visualize cell type distribution for each cell group; (7) GO enrichment analysis: Gene ontology enrichment analysis to reveal the activated/inhibited GO process for selected cell type and test/control groups. (8) Proliferation and EMT: Compute proliferation and EMT score for selected cell type (mainly used for cancer dataset); (9) Ligand & Receptor communication: Identify up-regulated ligands and receptors for each cell type and potential ligandreceptor interactions between different cell types given the selected test/control groups; (10) Dysfunctional signaling pathway: Identify dysfunctional cell-cell communication and signaling pathway between two cell types given the selected test/control groups; (11) Drug discovery: Identify possible drugs to inhibit the discovered cell-cell communication network. Note that the steps of pre-analysis (1)-(5) need to be done sequentially (indicated by blue color in the figure). All downstream analyses like (6)-(11) can be performed based on the interest after sc2MeNetDrug obtain the cell annotation results (indicated by orange color in the figure).

### The scRNA-seq data pre-analysis module

Recently, there have been many great scRNA-seq tools publicly available^24^ that integrate many aspects of analyses of scRNA-seq data. For example, Seurat^25^ in R and Scanpy^26^ in Python become very popular and be used as standard tools for analyzing scRNA-seq data. They include most of the common pipeline that is needed in the scRNA-seq data analysis like quality control, dimension reduction, cell clustering, differential gene expression analysis^22^, etc. However, one drawback of such tools is that they always require advanced knowledge in programming, which is not the case for many biomedical experts. Regarding this, sc2MeNetDrug implements^26, 27^the scRNA-seq pre-analysis module, which is a pipeline that includes quality control, normalization, imputation ^28^dimension reduction, clustering, gene feature visualization^24, 29^and cell type annotation^30^. The pre-analysis module is powered by Seurat and further adds many useful functions for easy processing of the scRNA-seq data. Most importantly, all methods are encapsulated into modules with user-friendly interfaces (**Fig 2a, 2c, 2d**), which make it easy for researchers to use even without programming skills.

**Figure 2:**
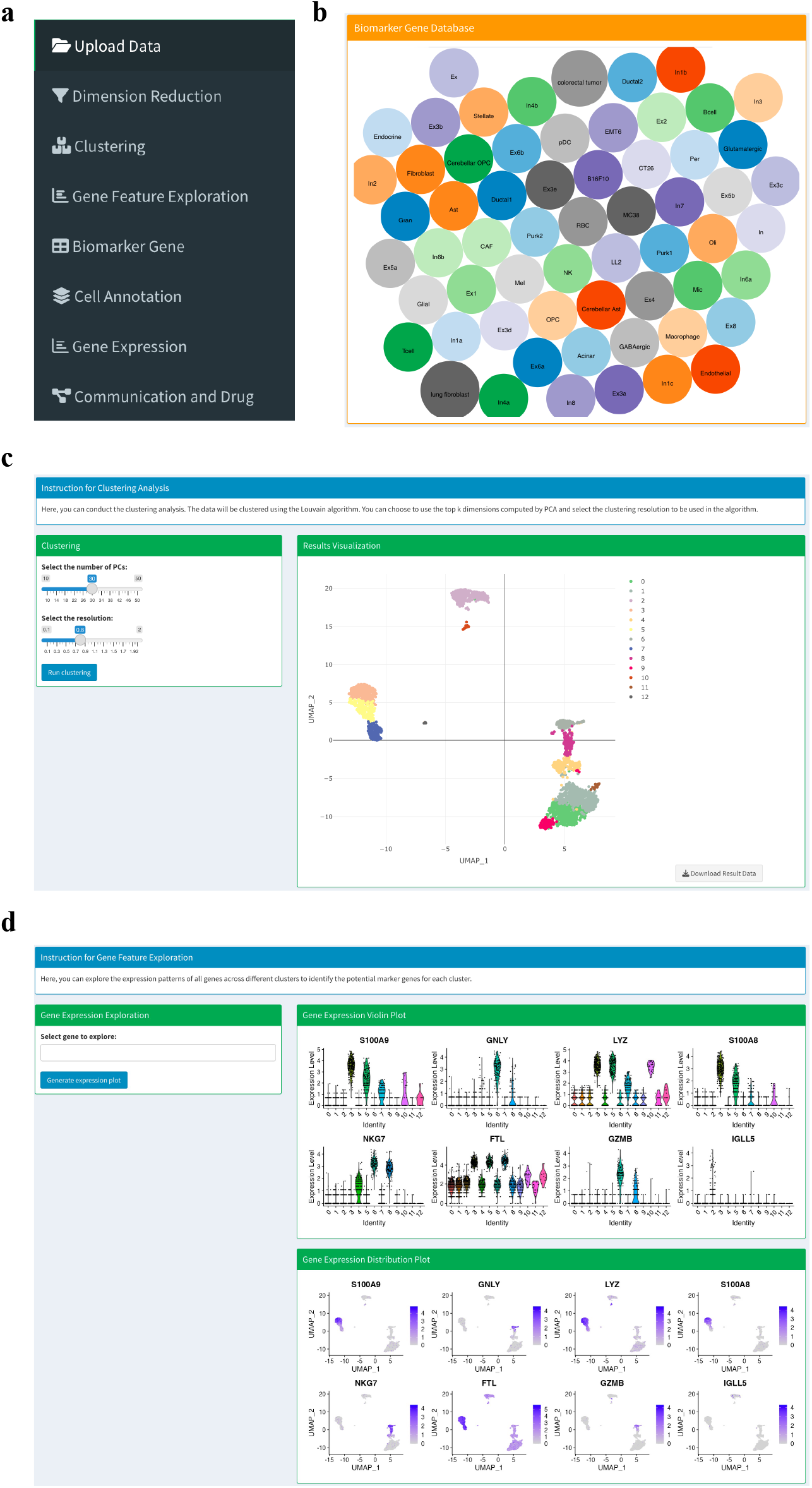
T**h**e **interface of some pre-analysis modules in sc2MeNetDrug.** (**a**) The menu bar of sc2MeNetDrug. (**b**) Biomarker gene database in sc2MeNetDrug. (**c**) The cell clustering section in sc2MeNetDrug. User can easily adjust parameters used in the algorithm. (**d**) The gene feature exploration section in sc2MeNetDrug used to identify biomarker genes.

In sc2MeNetDrug, both mice and human scRNA-seq data can be analyzed (mice gene symbols will be converted to corresponding human gene symbols). The quality control and data normalization will be computed automatically after the user uploads raw data. For pre-analysis, users can then do the dimension reduction and clustering analysis in order to perform cell annotation. Important parameters for each analysis can be adjusted directly in the app (**Fig 2c**). A large set of biomarker genes were collected^12,31–33^ to support different research projects, like cancer cells, immune cells, AD neuron cells (see **Fig 2b**). We will keep updating the marker gene sets. Moreover, we provided a function to enable users to upload new or user-defined marker gene sets. Once the users have decided on the final cell type candidates and their corresponding biomarker genes, the annotation classifiers based on these selected cell types and corresponding marker gene sets will be built automatically for the cell type annotation analysis. Also, the distribution (percentage) of individual cell types in each sample will be displayed, and the Epithelial–mesenchymal transition (EMT) and PRO (proliferation) scores of each sample can be calculated. Using the sc2MeNetDrug, we process and analyze the PDAC cohort from scratch. We can see sc2MeNetDrug successfully annotates each cell type in the dataset (**Fig 3a**). From the cell population result, we can verify the correctness of the annotation result, as Ductal 2 cells only exist in tumor groups (patients T1-T24) (**Fig 3b**). Ductal 2 cells are a well-known cell type related to tumor growth^33^. We also plot the EMT-PRO score for one tumor patient. We can see this patient has a high EMT score, which may indicate the high activity level of the metastatic expansion and the generation of tumor cells (**Fig 3b**).

**Figure 3:**
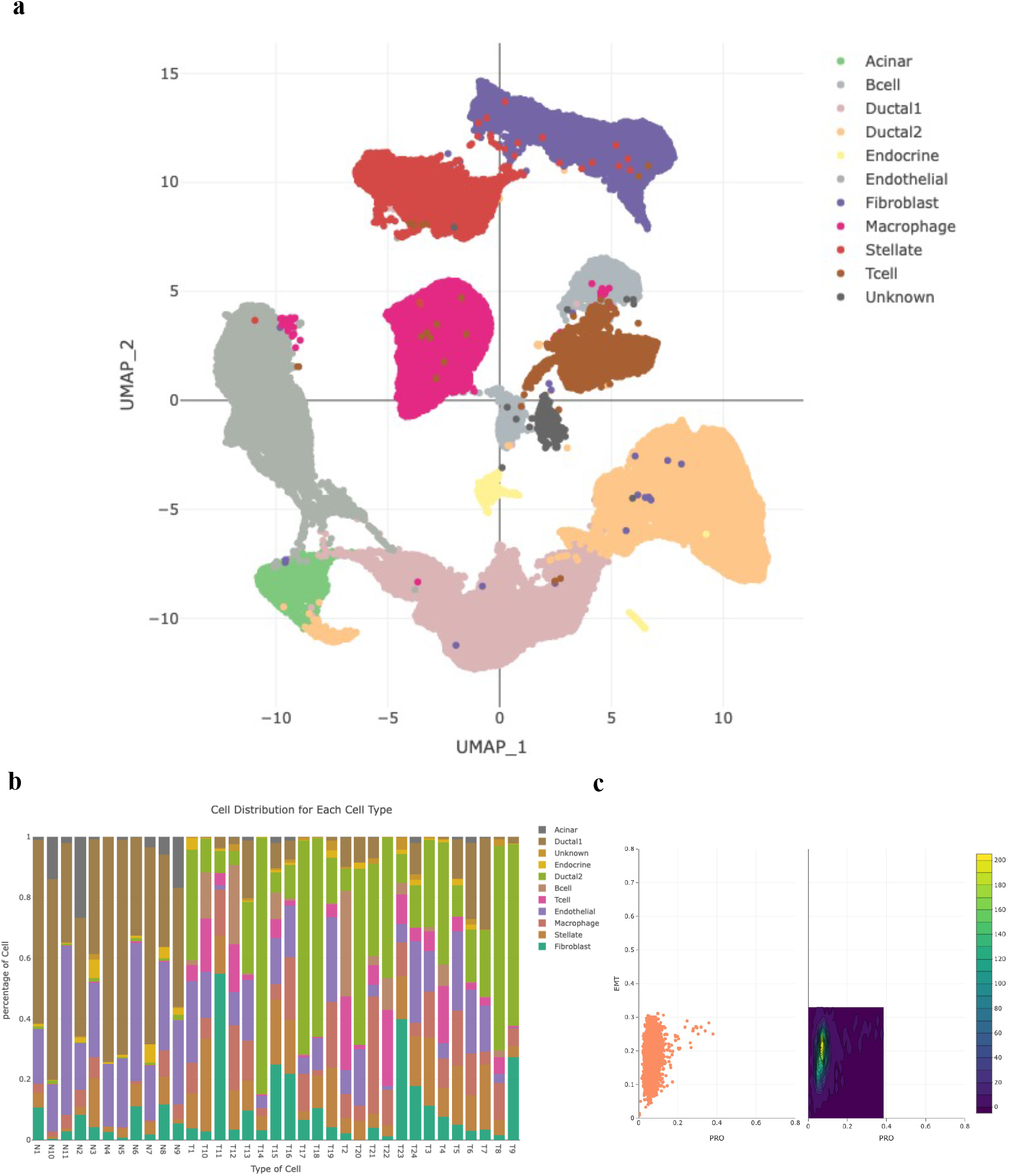
Analysis result of PDAC cancer using sc2MeNetDrug. (**a**) Cell annotation results. Sc2MeNetDrug clearly identify the cell type of each cluster. The cluster with negative enrichment score on all cell types is labeled with unknown to avoid noise. (**b**) Cell distribution in each group, which will be automatically plot after cell annotation. Tumor groups (T1-T24) have high population of Ductal 2 cell. Instead, normal groups only have Ductal1 cell. (**c**) EMT-PRO score of Fibroblast cells in one tumor patient.

### Identifying key biomarkers in Alzheimer’s disease using the iCSC module

Uncovering the dysfunctional signaling pathways within individual cell types, and cell-cell signaling communications, as novel therapeutic targets, are the highly needed functions. The SC2MeNetDrug provided functions to facilitate the pathway and network analysis. Specifically, after the cell type annotation, the differentially expressed genes in each cell type between two different experimental conditions, for example, the immunotherapy responder vs. non-responder, male vs. female, or tumor cells co-cultured with macrophage vs. no macrophages, can be calculated. A function was developed to enable the selection of samples and conditions of interest for the differential gene expression analysis. Based on the differentially expressed genes within individual cell types, gene ontology (GO) enrichment analysis can be identified. Further ligand-receptor interaction, activated signaling pathway, and cell-cell communication among two cell types can be computed accordingly.

To illustrate the functionality of sc2MeNetDrug for pathway and network analysis, we apply the sc2MeNetDrug on two AD cohorts, one from mice^14^ and another from human^12^. The mice cohort collected single-cell data from normal mice (TE3, TE4), mice with tau pathology and APOE3/APOE4 marker genes (TAFE3_oil and TAFE4_oil respectively), and mice with APOE3/APOE4 knock-out (TAFE3_tam and TAFE4_tam respectively). The human cohort was collected from 48 patients with 24 patients, 15 patients, and 9 patients classified as having No AD pathology, AD early-stage, and AD late-stage respectively. For the mice cohort, we apply the sc2MeNetDrug and use the tool to do the pre-analysis to obtain the cell annotation results. We can see sc2MeNetDrug clearly identifies marker genes for each cluster in the cohort (**Fig 4a, 4b**) and annotates each cell type in the dataset (**Fig 4c**). For the human cohort, we directly use the cell annotation result from the original source in order to validate the functionality of the downstream analysis part in sc2MeNetDrug. For mice dataset, we use TE4 as the control group and TAFE4_oil as the test group. For the human dataset, we use No AD pathology as the group of control and late-stage as the group of test. Next, we conduct the GO enrichment analysis on both mice and human cohorts using the sc2MeNetDrug to compare the difference of neurons between normal and AD pathology. We further conduct the KEGG pathway enrichment analysis company with the GO enrichment analysis to provide a complete view (**Fig 4e**). From the result of the mice dataset, we have the following observations: First, the neuron autophagy and degeneration-related processes are highly activated in neurons with AD pathology, like *Pathway of neuron degeneration – multiple disease*, *Apoptosis* in KEGG results and *Neuron apoptotic process, Autophagy* in GO results. Autophagy is a lysosome-dependent, homeostatic process, in which organelles and proteins are degraded and recycled into energy. Autophagy has been linked to Alzheimer’s disease pathogenesis through its merger with the endosomal-lysosomal system, which has been shown to play a role in the formation of the latter amyloid-β plaques^34^. In the prediction of sc2MeNetDrug, we also identified several important genes related to autophagy (**Fig 4f**). Some have already been shown to be related to neuron degeneration and autophagy in AD like *FAIM2*^35^, *BCL2*^36^, and *PRNP*^37^. One hypothesis is that irregular autophagy stimulation results in increased amyloid-β production^38^. Our result also identified the highly up-regulated GO term *Positive regulation of amyloid-beta formation*, which further supports it. Secondly, neuron inflammation is prevalent in AD pathology. Numerous studies have shown that inflammation is highly activated and plays a key role in the progress of AD^12,39–41^. Our results further verify this claim. We can see that *neuroinflammatory response*, *cytokine-mediated signaling pathway* are all up-regulated GO terms discovered by sc2MeNetDrug. This result aligns with the previous studies and further confirms that the existence of APOE4 in the astrocyte stimulates the inflammatory response. Inflammation-related genes are also identified (**Fig 4f**) like *TREM2*, *CLU*, and *ADCY1*. We further use sc2MeNetDrug to compute the activated KEGG signaling pathway network for excitatory neurons using mice cohort (**Fig S1**). The result points out additional pathways like *Estrogen signaling pathway*, *HIF-1 signaling pathway*, and *MAPK signaling pathway* that are activated in the AD neurons. Some works have pointed out that the dysfunction of the estrogen signaling pathway also contributes to the production of amyloid-beta and the progress of AD^42^. Particularly, gene CTSD is Highly expressed in neurons, suggesting its center role in producing amyloid precursor (APP) and tau. MAPK signaling pathway regulates a variety of cellular activities including proliferation, differentiation, survival, and death. Some studies report that Amyloid-beta-induced activation of p38 MAPK and NFkB signaling can result in upregulation of proinflammatory gene transcription and cause neuronal death^43^.

**Figure 4:**
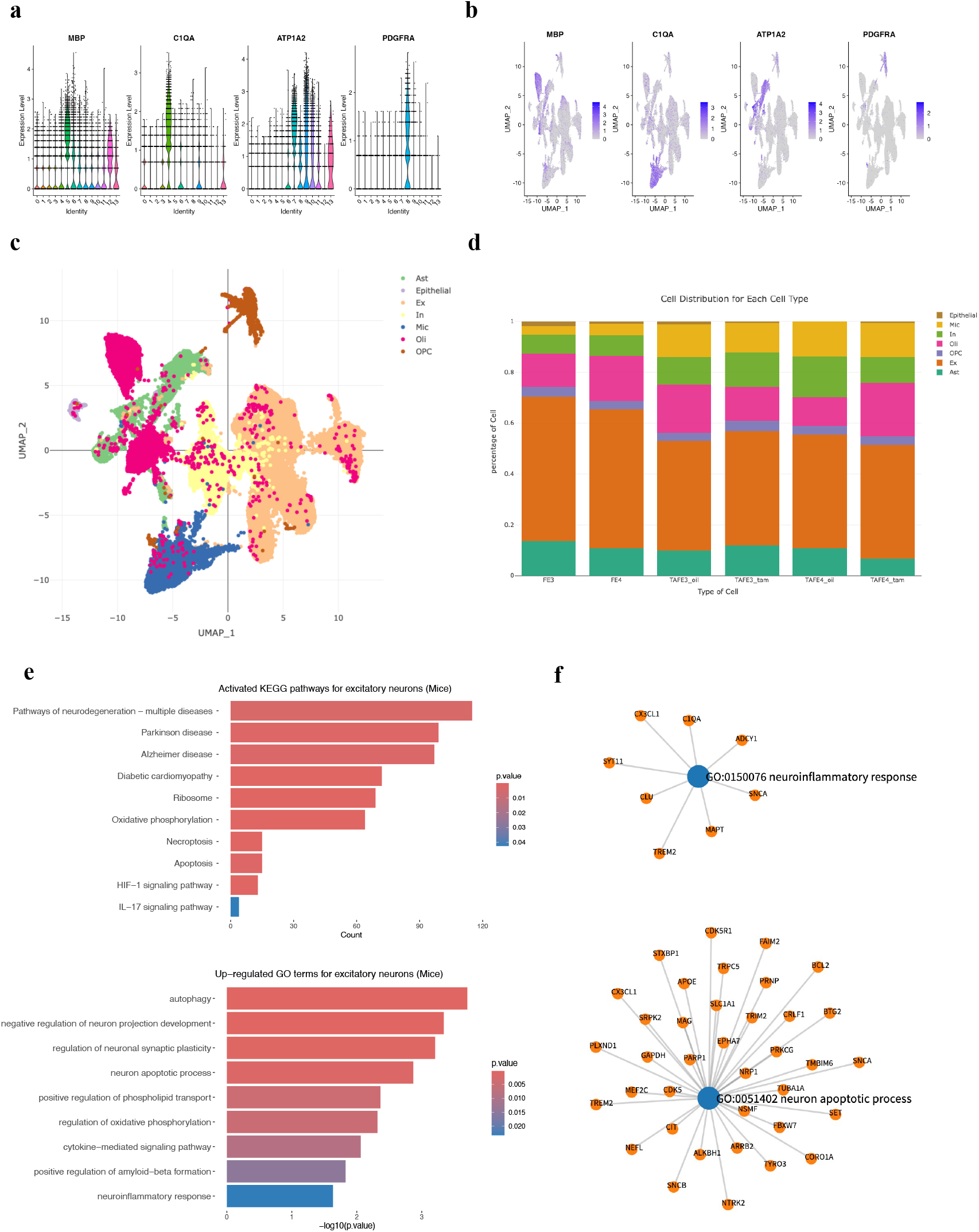
**Cell annotation and pathway analysis result of AD (mice) dataset using sc2MeNetDrug**. (**a-b**) The expressive pattern of selected maker genes. These genes are highly expressed in some cluster and can be used as biomarker genes for cell annotation. (**c**) The cell annotation result output from sc2MeNetDrug. (**d**) The cell distribution for each group. (**e**) The KEGG and GO enrichment analysis results for AD pathology. The color indicate the p-value of pathways and count (KEGG pathway) indicate the number of genes that are activated in the pathway. The log fold-change threshold is set as 0.08 and p-value threshold is set as 0.05. (**f**) Expressed genes correlated to GO terms GO:0150076 neuroinflammatory response and GO:0051402 neuron apoptotic process for AD pathology output by sc2MeNetDrug. The log-fold change threshold is set as 0.08.

To further investigate the signaling pathway and disease mechanism of AD, we apply sc2MeNetDrug on two cohorts to analyze both the ligand-receptor interaction and inter-cell communication patterns. First, we use sc2MeNetDrug to compute the up-regulated ligands and receptors for both mice and human cohorts (**Fig 5a**). For excitatory neurons, *PCSK1N*, *ALDOA*, *CLU*, *PRNP*, and *LINGO1* are highly up-regulated in both mice and human cohorts. For astrocytes, *PTGDS* and *CLU* are activated in mice and humans commonly. For microglia, we found that *APOE* is highly up-regulated in both mice and human cohorts, even though the *APOE* is normally expressed in astrocytes. This may indicate the APOE in the microglia may also be a critical factor for the development of AD pathology. Besides that, *RPS19*, *SPP1* are also highly expressed in AD pathology. The differential gene analysis result further confirms the discovery from sc2MeNetDrug (**Fig 5c, 5d**).

**Figure 5:**
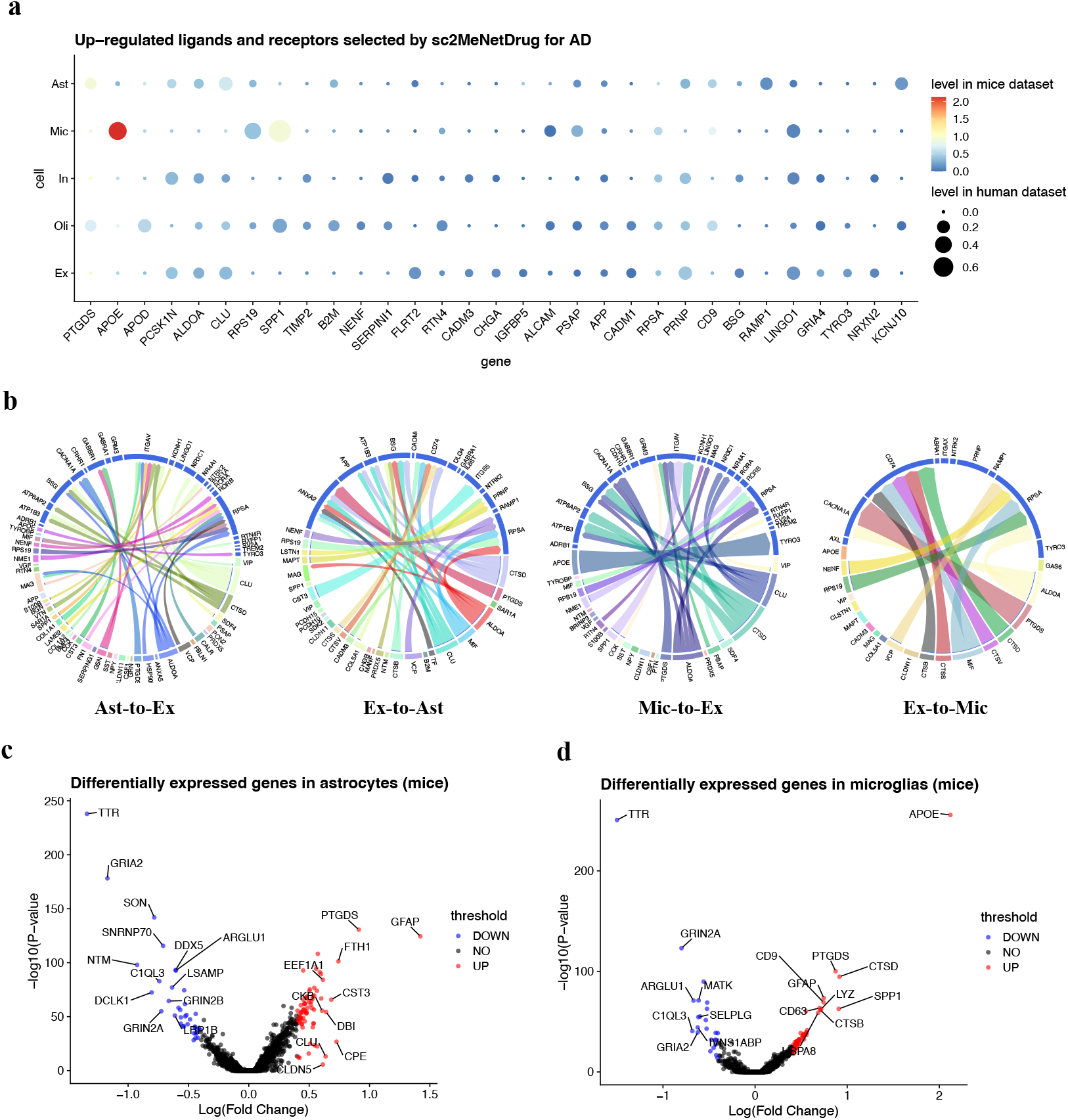
**Ligand-receptor analysis results on AD cohorts**. (**a**) Up-regulated ligands and receptors discovered by sc2MeNetDrug on two AD cohorts. Only genes that appear in the results on both mice and human cohorts are selected. The log-fold-change threshold is set as 0.08 and p value threshold is set as 0.05. The size of the dot indicates the fold log-fold-change level in the human cohort and the color indicate the level in the mice cohort. (**b**) ligand-receptor interaction discovered by sc2MeNetDrug on mice cohort. Only ligands and receptors with log-fold-change great than 0.08 were selected. (**c-d**) Differentially expressed genes results using Seurat.

Finally, we use sc2MeNetDrug to discover the up-regulated-ligand to up-regulated-receptor interaction and cell-cell communication networks among excitatory neurons, astrocytes, and microglia (**Fig 5b**, **Fig 6a**). The results strengthen the understanding of AD development and neuron change. First, *COL1A1, COL6A1, COL16A1* are differentially expressed in astrocytes of AD pathology. It all connect to the common receptor ITGAV in neuron cell and further connect to genes like *PIK3CA, ACTG1, and ACTB*. The collagen gene family serves to mediate cell attachment and maintains the integrity of the extracellular matrix (ECM). It has been reported that there are significant changes in ECM during the early stages of Alzheimer’s disease^44^ and also associated with amyloid plaque production^45^. *ACTG1* and *ACTB* are actin proteins, which are highly related to actin cytoskeleton and spine shaping in the brain. The abnormal expression of actin-related genes can cause synaptic plasticity and failure, which are one of the major remarkers of AD. Our finding may suggest that the activity of collagen genes in astrocytes may trigger the abnormal activity of actin-related genes and thus contribute to the development of AD pathology. Further, our identified cell-cell communication network contains many genes in the activated pathways, like Apoptosis, HIF-1 signaling pathway, Necroptosis, and neuroinflammatory response. The results further highlight the significance of these pathways and processes in the progress of AD pathology. Meanwhile, these findings are consistent with results from the pathway enrichment analysis, further validating our iCSC module.

**Figure 6:**
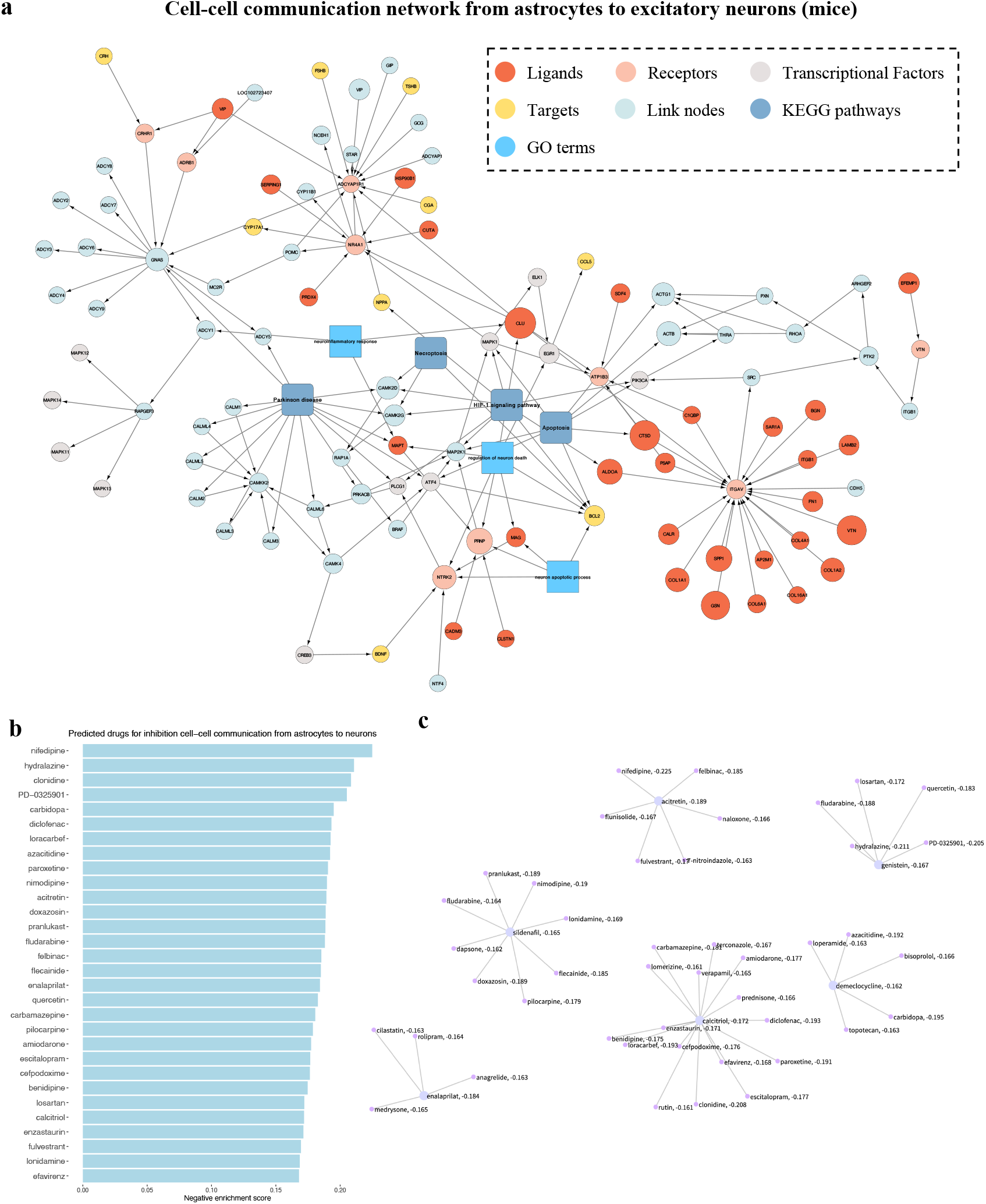
Cell-cell communication result from astrocytes to excitatory neurons and corresponding drug discovering result (mice cohort) using sc2MeNetDrug. (**a**) Cell-cell communication result with several important pathways appended. The KEGG database was used and the log-fold-change and p-value threshold are set as 0.08 and 0.05 respectively. (**b**) Signature drug discovering result. Only top 30 drugs are displayed. The higher the negative enrichment score, the higher chance the drug can be used to inhibit the corresponding dysfunctional network. (**c**) Drug clustering result based on SMILE finger print to identify structure similar drug clusters among all top drugs.

### Predict drugs inhibiting signaling communications using the dCSC model

To identify drugs that can potentially inhibit the down-stream signaling pathways, the computational model, the ***dCSC*** model was developed, which is designed to integrate the down-stream signaling network, drug-target interaction (derived from DrugBank^46^ database), and reverse gene signature data available from the connectivity map (CMAP) database^47^. It is relatively straightforward to identify drugs with targets on the downstream signaling network based on the drug-target information. For the CMAP data^48^, the uncovered cell-cell communication signaling network will be used as a signature of the Gene set enrichment analysis (GSEA) to identify drugs that can potentially inhibit the expression of genes in the network. To further understand the relationship of the selected drugs, drug clustering based on chemical structures was conducted to identify therapeutics with similar targets or mechanisms of action. To demonstrate the functionality and effectiveness of the dCSC module in sc2MeNetDrug, we perform the analysis using the AD mice dataset. Specifically, we compute the signature drug discovering analysis using connectivity map data upon the cell-cell communication network from astrocytes to neurons. The discovered top 30 drugs can be found in **Fig 6b**. The drug clustering result for all top drugs can be found in **Fig 6c**.

Among all the top drugs, Nifedipine achieved the best enrichment score. Nifedipine is a dihydropyridine calcium channel blocker indicated for the management of several subtypes of angina pectoris, and hypertension. Nifedipine is primarily used for reducing blood pressure and increasing oxygen supply to the heart. However, results have shown that calcium channel blockers like Nifedipine or Nimodipine (also shown in the top drugs) led to a significant decrease in the level of amyloid beta peptide, with no significant decrease in cell viability^49^.

The second drug Hydralazine, which is a smooth muscle relaxant, also shows potential for being an effective drug for reducing lipid oxidative damage and amyloid beta folding^50^. Interestingly, another drug PD-0325901 from the same cluster, has been shown to be effective in inhibiting MAPK/ERK kinase which prevents activation of the mitogen-activated protein kinase (MAPK)^51^. As we discussed above, MAPK signaling pathway is dysfunctional in AD pathology. The inhibition of MAPK pathway can provide neuronal protection from the amyloid beta burden by increasing autophagic lysosomal activity, suggesting a synergic effect along with Hydralazine that is worth for further investigation. Another type of drug mainly targets neuronal inflammation or neuronal disorders like carbidopa, Diclofenac, and paroxetine. Relevant research already pointed out its potential functionality in reducing inflammation and pathology of AD^52–54^. Together, our results provide several drugs that could be effective in reducing the pathology of AD from different levels. It further valid the performance of sc2MeNetDrug.

## Discussion

Single cell RNA sequencing (scRNA-seq) is a powerful technology to investigate the transcriptional programs in stromal, immune, and tumor cells or neuron cells within tumor or brain microenvironment (ME) or niche. Cell-cell interactions and communications within ME play important roles in disease progression and immunotherapy response and are novel and critical therapeutic targets. However, it is challenging, for many researchers without solid training in computational data analysis and scRNA-seq data analysis, because the data analysis pipelines usually consist of diverse and complex analysis modules, and the integrative analysis of diverse and heterogenous external data resources. There is a lack of easy-use tools with complete and integrative computational modules for uncovering cell-cell communications of ME and predict the potentially effective drugs to inhibit the communications, although many tools of scRNA-seq analysis have been developed to investigate the heterogeneity and sub-populations of cells. In this study, we developed a novel, open-source computational tool, SC2MeNetDrug (https://fuhaililab.github.io/sc2MeNetDrug/) to address these challenges. Specifically, the advantages of the tool are as follows. First, it is a tool specifically designed for scRNA-seq data analysis to identify cell types within MEs, uncover the dysfunctional signaling pathways within individual cell types, inter-cell signaling communications, and predict effective drugs that can potentially disrupt cell-cell signaling communications. Second, the analysis modules in the analysis pipelines were separated with pre-designed interfaces. Users can develop and update novel data analysis modules, and easily replace the updated modules back to the data analysis pipeline. In another word, users or scientists with different expertise can conveniently replace user-specific data analysis modules just by following the input and output of individual modules, like network inference, cell type identification, cell clustering, drug prediction, in the data analysis pipeline. Third, it provides a user-friendly graphical user interface (GUI), encapsulating the data analysis modules, which requires no coding and programming and can facilitate the scRNA-seq data analysis in an interactive manner.

## Conclusion

In this study, we developed a novel open-source tool, SC2MeNetDrug (https://fuhaililab.github.io/sc2MeNetDrug/), which is specifically designed, with user-friendly GUI for interactive scRNA-seq data analysis for the purpose of uncovering cell-cell communications of ME, and predicting the potentially effective drugs to perturb the cell-cell communications within disease ME.

## Methodology

### Data sets of case studies

The data of PDAC was downloaded from Genome Sequence Archive under project PRJCA001063^16^. There are a total of 57530 cell samples and 24003 genes. The data was generated from 24 PDAC tumor samples and 11 control, untreated pancreas samples. The data of Alzheimer’s disease of human^12^ was downloaded from Synapse website. The DOI for the dataset is 10.7303/syn18485175. The data was generated from 48 patients, with 24 individuals presenting no or very low AD pathologies and the rest of 24 individuals exhibiting clear AD pathologies. There are a total of 75060 cell samples and 17926 genes. The data on Alzheimer’s disease in mice^14^ was obtained from the Gene Expression Omnibus (GEO) database with accession number GSE164507. There are a total of 96252 cell samples and 33457 genes.

### Quality control

Quality control is done in several steps. Initially, cells with a detected gene count of less than 200 or more than 7500 are removed. Subsequently, cells with abnormal mitochondrial gene expression (cells with > 10% mitochondrial counts) are also eliminated. Finally, if the dataset is from mice, we collect a database for converting all mic gene symbols in the dataset to the human gene symbol. All genes that don’t exist in the database will be removed.

### Normalization

The normalization and variance stabilization of scRNA-seq data in sc2MeNetDrug is done by the regularized negative binomial regression^55^. With this method, there is no need for heuristic steps including pseudocount addition or log-transformation and improves common downstream tasks. The method is implemented using *SCTtransform* function with *method=glmGamPoi* in Seurat package^56^.^56^

### Imputation

Imputation is done by the *runALRA* function in the Seurat package with default parameters. The method^28^ is to compute the K-rank approximation to A_norm and adjust it according to the error distribution learned from the negative values.

### Dimension reduction

Dimension reduction analysis in sc2MeNetDrug involves several steps. First, select the top 3000 variable genes across the dataset. To identify these genes, local polynomial regression fits the relationship between log variance and log mean. Subsequently, gene expression values are standardized using the observed mean and the expected variance (determined by the fitted line). The variance of gene expression is calculated on the standardized values after clipping. This procedure is automatically executed by the *SCTransform* function in Seurat package. Next, Principal Components Analysis (PCA) is applied to these 3000 variable genes. These genes are then projected into 50 dimensions in order served as 50 different principal components (PCs). This is implemented using the *RunPCA* function in Seurat package. Finally, the UMAP method will be used on the first *x* PCs selected by the user (range from 10 to 50). This is implemented using the *RunUMAP* function in Seurat package.^57^

### Cell clustering

For clustering analysis of scRNA-seq data, the application follows these steps. First, it computes a low-dimensional representation for each cell using PCA. This is done during the dimension reduction analysis. Next, to identify the neighbors of each cell, the clustering analysis applies the K-Nearest-Neighbor (KNN) algorithm to the results of the PCA analysis. Users can choose the number of principal components to use (range from 10 to 50) when performing the KNN algorithm. This step is implemented using *FindNeighbors* function in Seurat package. Finally, the clustering analysis applies the Louvain algorithm to the results of the KNN algorithm to compute the final clustering results for each cell. Users can also choose the resolution of the clustering, with options ranging from 0.1 to 2. A lower resolution result in fewer clusters in the final results, while a higher resolution produces more clusters. This step is implemented using *FindClusters* function in Seurat package.

### Gene Feature Exploration

In gene feature exploration section, user can select any gene exists in the dataset, then sc2MeNetDrug will generate two plots for each gene. The first plot is the expression distribution violin plot, which plot the expression distribution of the selected gene in each cluster. The second plot is the scatter plot to indicate the expression of the gene in each cell. A deeper color means a higher expression. These two types of plots are drawn using *VlnPlot* and *FeaturePlot* function in Seurat package.

### Biomarker gene sets

In total, we collected 56 cell type and biomarker genes from several sources^12,31–33^. The biomarker gene database is displayed in a table with a cell marker gene manner, where each row in the table has two columns with the first column indicating the name of cell type and the second column indicate the corresponding gene symbol. If there is more than one biomarker genes for one cell type, each gene will be displayed in one row (the cell type name will be copied for each row, see **Fig 7a**). We also specified classical cell type sets for Alzheimer’s disease and Pancreatic Cancer based on published articles^12,33^. The user could easily select these cell types by clicking the corresponding button. We also provide the user with the ability to modify and add their own marker genes for better analysis; the user can add, delete and modify existing marker gene tables. To delete an existing cell-marker gene pair, user can do it by first clicking the corresponding row in the marker gene table. Then, click the “Delete selected gene” button to delete this row (see **Fig 7b**). To add a new cell-marker gene pair, users can use the bottom right panel in the biomarker gene section (**Fig 7c**). Begin by entering the cell type name in the first input box. Next, select the marker genes you wish to add in the second input box. Users can choose multiple genes simultaneously. Lastly, click “add new gene” to incorporate them into the marker gene database. Finally, to modify an existing cell-marker gene pair, users can directly double-click the table cell in the marker gene table to modify it.

**Figure 7:**
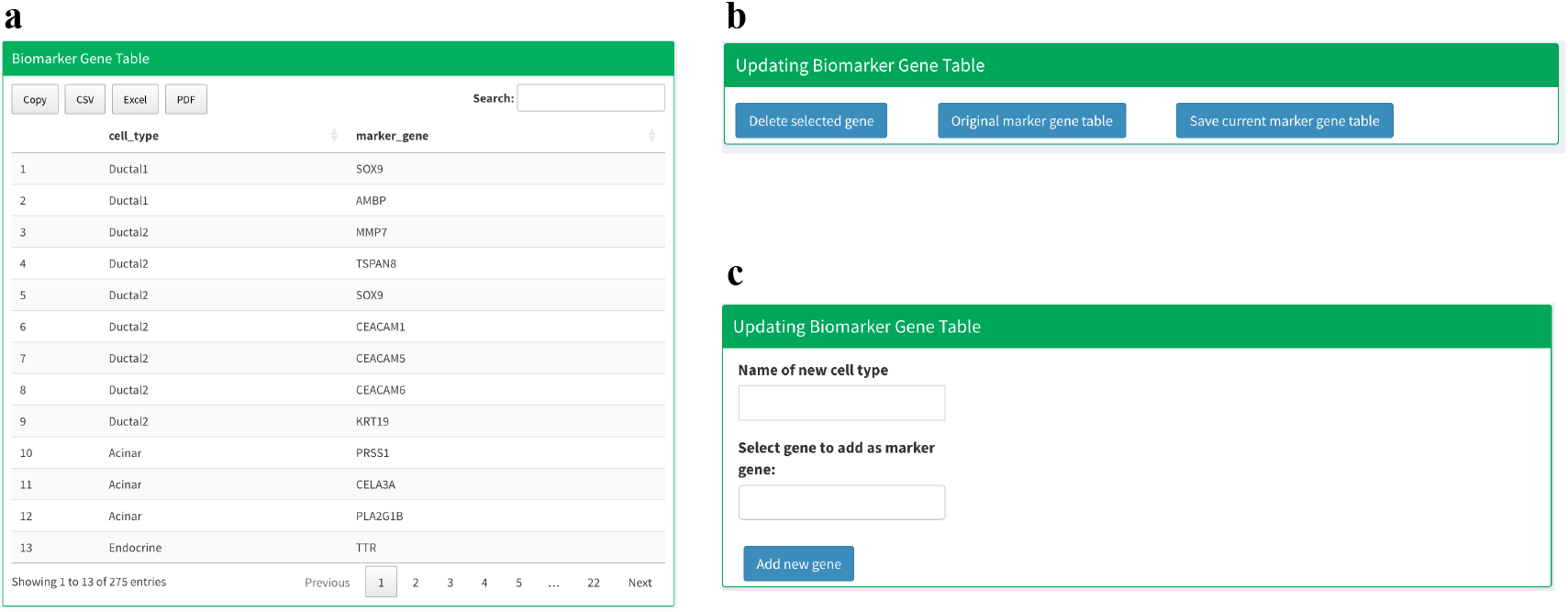
Panel shortcut for modifying biomarker gene database in sc2MeNetDrug. (a) Biomarker gene table display all existing cell-marker gene pairs. **(b)** The panel used to delete an existing cellmarker gene pair (“Delete selected gene” button), reset the biomarker gene database (“Original marker gene table” button), and save the modified marker gene database for future use (“Save current marker gene table” button). **(c)** Panel used to add new cell-marker gene pair into current biomarker gene table.

### Cell type annotation

The cell annotation is done in two steps. In the first step, the Gene Set Enrichment Analysis (GSEA)^59^ is applied to annotate cell types for every cluster. First, users should select candidate cell types and corresponding marker genes in the Biomarker gene section. Then, for every cluster, the application computes log fold change for cluster N by:

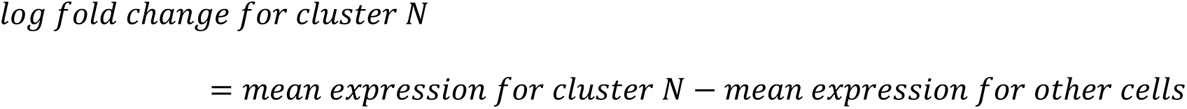

Then we rank the genes based on fold change and calculate the enrichment score of marker gene sets for every cell type the user selected. Finally, the cell type with the largest enrichment score will be selected as the type of this group. However, if none of the cell types have a positive enrichment score, the cluster will be annotated as unknown. In the second step, sc2MeNetDrug introduces a manual label correction panel such that the user can manually modify the cell annotation result obtained from the first step.

### Cell distribution plots

Once users obtain classification results or upload gene expression data, the application can calculate the percentage of each cell type in each sample group. If users don’t provide sample group information, the application will simply calculate the percentage of each cell type in the whole dataset.

### Epithelial-mesenchymal transition (EMT) and proliferation (PRO) analysis

EMT-PRO analysis in SC2NetDrug was analyzed by computing mean expressions for the selected design and cell type of EMT and PRO-related genes. The HALLMARK_EPITHELIAL_MESENCHYMAL_TRANSITION database was chosen for EMT-related marker genes and the HALLMARK_E2F_TARGETS database was chosen for PRO-related marker genes. After users select the design and cell type, min-max normalization is used to normalize the whole dataset based on the genes. Then the intersecting genes in the EMT and PRO-related marker gene sets are selected and a mean score of all EMT and PRO-related genes are calculated and labeled as the EMT and PRO scores, respectively.

### Ligands and receptors data resources

We collected ligand-receptor data from several sources: (1) Database of Ligand-Receptor Partners (DLRP)^60^ with 175 unique ligands, 133 unique receptors, and 470 unique interactions (2) Ligand-receptor interaction data sources in NicheNet^23^ with 1737 unique ligands, 1925 unique receptors and 12659 unique interactions (3) cell-cell interactions database in baderLab. We selected all the proteins to be annotated as ligands named “Ligand” or “ECM/Ligand” and all the proteins to be annotated as receptors named “Receptor” or “ECM/Receptor”. Then we selected all the interactions including the chosen ligands and receptors. There are 1104 unique ligands, 924 unique receptors, and 16833 unique interactions. In total, there are 1424 unique ligands, 1214 unique receptors, and 27291 unique interactions.

### Ligand-receptor mediated signaling interactions (Upstream Network)

Upstream network analysis is used to discover up-regulated ligands, receptors, and potential ligand-receptor signaling interactions. First, users need to specify the log fold change threshold, p value threshold, and the group or design the user wants to analyze. The up-regulated ligands and receptors are discovered using the following steps. First, the differential expression genes are calculated based on two tests, the first being the Wilcoxon rank sum test and the second being the Likelihood-ratio test^61^. The genes that have log fold changes larger than the threshold and adjusted p-values (from the two tests) less than the threshold will be selected as differentially expressed genes. The test is done by the *FindMarkers* function in the Seurat package with parameters set as *test.use=wilcox* and *test.use=bimod* for the two tests respectively. After the differentially expressed genes for all cell types in the dataset designed by the user are identified, the ligands and receptors are found by searching for all differentially expressed genes in our ligands-receptors database. Finally, upstream interaction networks are generated by searching for all the discovered ligand-receptor interactions in our ligands-receptors database. To be specific, four networks will be generated: the up-regulated ligands to expressed receptors network, expressed ligands to up-regulated receptors network, up-regulated ligands to up-regulated receptors network, and the combined network. Up-regulated ligands and receptors are ligands and receptors that have log fold changes and adjusted p-values for two tests that satisfy the user’s settings. Expressed ligands and receptors are ligands and receptors that have log fold changes larger than 0. The combined network is then combined with the up-regulated and the expressed ligands and receptors.

### Gene ontology (GO) term enrichment analysis

To obtain the gene-gene ontology (GO)^62^ term information, the R libraries, org.Hs.eg.db and GO.db were used. The Fisher’s exact test was used to identify the statistically activated/enriched GOs based on the up-regulated genes and the genes in each GO term.

### Inter-Cell Communication (Downstream Network) Analysis

The inter-cell communication analysis in SC2NetDrug is done by several steps. First, differential genes in each cell type are discovered using the Wilcoxon rank sum test and the Likelihood-ratio test^63^. The genes that have log fold changes larger than the threshold and adjusted p-values (for both tests) less than the threshold will be selected as differentially expressed genes. The tests are done by the *FindMarkers* function in the Seurat package with parameters set as *test.use=wilcox* and *test.use=bimod* for the two tests, respectively. Next, ligands, receptors and transcript factors are discovered using the ligands-receptors interaction database and the transcript factor-target interaction database.

To uncover the down-stream signaling of ligand-receptor of interest, a computational model, ***iCSC*** (**i**nter-**<u>c</u>**ell **<u>s</u>**ignaling **<u>c</u>**ommunication discovery using **sc**RNA-seq), was developed. Specifically, 2 background signaling resources were used: KEGG^64^ signaling pathways (curated) and STRING^65^ (general protein-protein interactions). For KEGG signaling pathways, the shortest paths starting from the given receptors to all the target genes (without out-signaling) were identified, denoted as 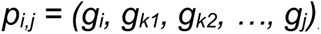, where *g_i_* is the receptor, *g_j_* is the target gene, and *g_km_*, m=1, 2, …, are the genes on the shortest paths between *g_i_* and *g_j_* on the KEGG signaling pathways. Then an activation score for each path, *p_i,j_*, was defined as: 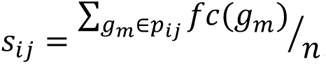, where *fc(.)* is the fold change calculator, and n is the number of genes on the signaling path. Then signaling paths with activation scores greater than a given threshold will be selected to generate the inter-cell communication network of the ligand-receptor of interest.

For STRING background signaling network, there are much more genes (nodes) and interactions (edges) than KEGG signaling. Thus, the above model for KEGG does not work for STRING. Herein, we proposed a novel down-stream signaling network discovery model. Specifically, let 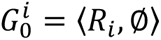 denote the initialized down-stream signaling network of receptor R*_i_*. The update of the down-stream signaling is defined as: 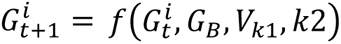, where 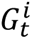 and *G_B_* is the current down-stream and background (STRING) signaling networks respectively. The edge, *e_ij_* (protein interactions between *g_i_* and *g_j_*) of background signaling network, *G_B_*, is weighted as: 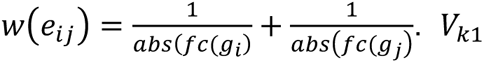. is a vector including k1 candidate genes (based on the absolute fold change in the decreasing order) to be investigated and added to the down-stream signaling network. For any gene, 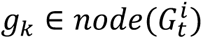, the shortest paths from *g_k_* to the k1 candidate genes in *V_k_*_1_, will be calculated. Then, an activation score for each path, *p_k,j_*, was defined as: 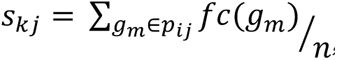, where *fc(.)* is the fold change calculator and n is the number of genes on the signaling path. If *n* > *k*2, the signaling paths will be discarded. In another word, the parameters k1 and k2 decide the search width and depth. Finally, the signaling path with highest activation score will be added to the down-stream signaling network. The process will be conducted iteratively until it reaches a network size limit, e.g., N nodes. The down-stream signaling network is generated by combining the down-stream signaling networks of all receptors: 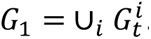.

### Drug-target information derived from DrugBank

We collected 6650 drugs from the drug bank database and corresponding target genes. After the down-stream signaling network is generated, the drugs for genes in the network is discovered by looking through each gene in the network and searching for drugs that target this gene in drug bank database.

### Connectivity Map data

Drug discovery based on signaling signatures using Connectivity Map data, which seeks to enable the discovery of functional connections between drugs, genes, and diseases through analysis of patterns induced by common gene-expression changes. Users can find CMAP data in National Center for Biotechnology Information database under dataset GSE92742. Before doing the analysis, users need to download corresponding data from the website and we provide the function to generate the drug rank matrix based on data.

### Drug discovery based on signaling signatures

The procedure of drug discovering is following: After the up-regulated genes for each cell group in the cell-cell communication part are obtained, the application will use GSEA and the drug rank matrix to discover potential drugs for each group. First, the application will calculate the enrichment score of up-regulated gene sets for each drug in each group. Then, the top K drugs with the lowest enrichment scores will be selected as potential drugs, where K is the number of top drugs selected by user.

### Drug clustering based on GSEA scores in CMAP

After the top drug is identified, Affinity Propagation Clustering^66^ will be used to cluster top drugs. First, a similarity matrix will be constructed for the top drugs. Given that the number of top drugs is K, the dimensions of the matrix will be K*K. The similarity score for drug *i* to drug *j* will be computed by the following process: select the top 150 up-regulated genes and top 150 down-regulated genes for drug *i* to use as the gene set. Then, compute the GSEA score for drug *j* using the drug rank matrix and the gene set from drug *i*. The enrichment score will be used as the similarity score for drug *i* to drug *j*. After the similarity matrix is constructed, it will be used to do AP clustering, which is done using the R package apcluster.

### Drug clustering based on chemical structures

To clustering drugs discovered by targets, we use the chemical structure of each drugs^67^. First, the SMILES information of drugs is used to generate drug object for each drug, this is done by *parse.smiles* function in rcdk R package. Next, the fingerprint of drug is computed using *get.fingerprint* function in fingerprint R package. Based on fingerprint of drugs, the similarity between drugs is computed using Tanimoto index. The formulation of Tanimoto index is follow:

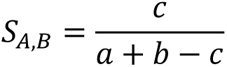

Where *S_AB_* is the similarity between drug *A* and drug *B*. *a* is number of bits in drug *A* and *b* is number of bits in drug *B*. *c* is number of bits in both two drugs. This is done by *fp.sim.matrix* function in R package fingerprint and set parameter method as *tanimoto*.

## Author contributions

FL conceived the project, JF and FL conducted the model, tool development and data analysis and wrote the manuscript. SPG, YB, AZ, DD, WH, RF revised the manuscript. SPG, TW, PP, LD, DD, WH, RF discussed the results.

## Acknowledgment

This study was partially supported by the Children’s Discovery Institute (CDI) M-II-2019-802, 1R01LM013902-01A1, R56AG065352, 1R21AG078799-01A1 to Li.

---

**Supplementary figure 1:**
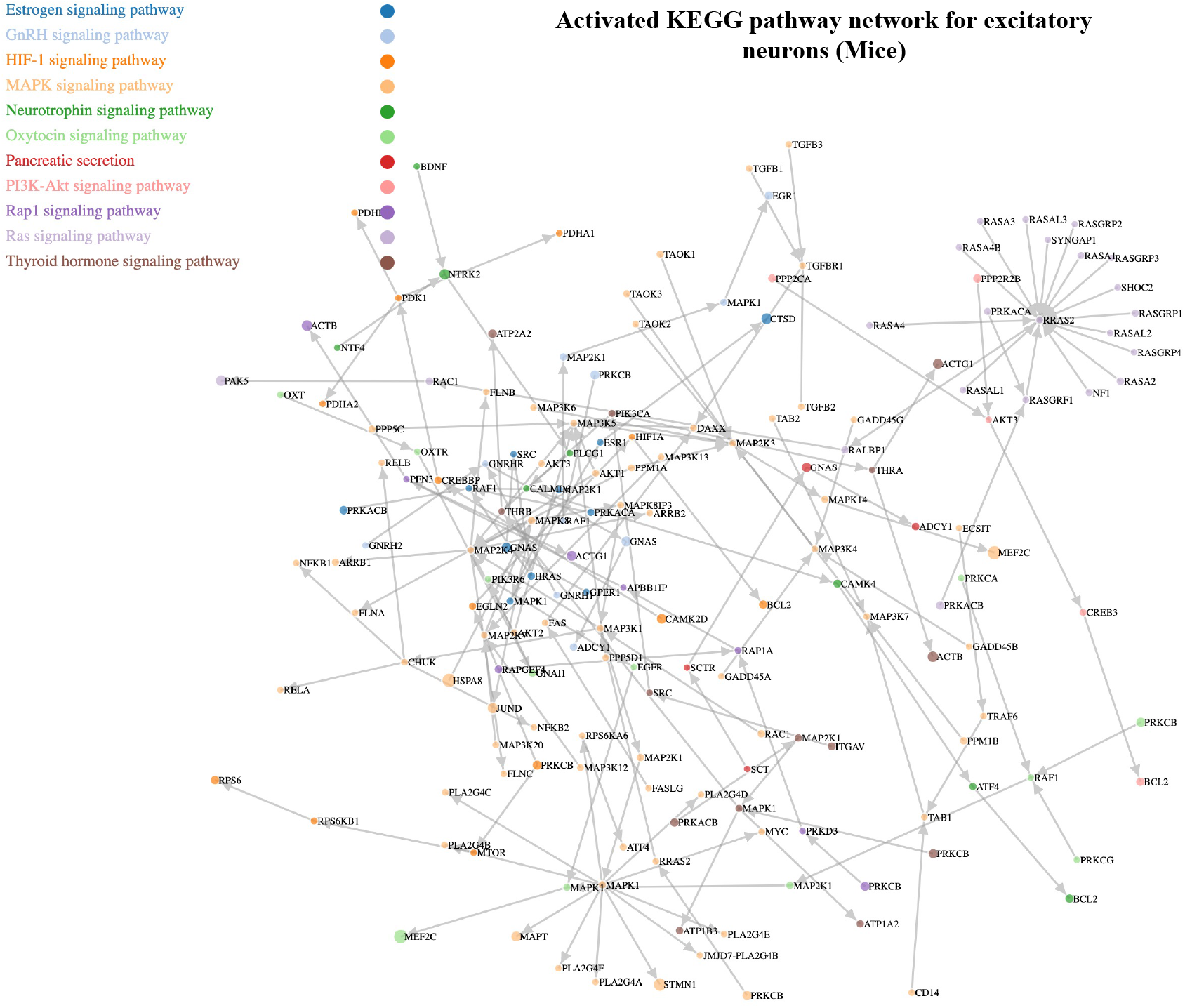
Activated KEGG pathway network of AD pathology for excitatory neurons
identified by sc2MeNetDrug (mice cohort).

